# The N terminus of adhesion G protein-coupled receptor GPR126/ADGRG6 as allosteric force integrator

**DOI:** 10.1101/2021.09.13.460127

**Authors:** Jakob Mitgau, Julius Franke, Camilla Schinner, Gabriele Stephan, Sandra Berndt, Dimitris G. Placantonakis, Hermann Kalwa, Volker Spindler, Caroline Wilde, Ines Liebscher

## Abstract

The adhesion G protein-coupled receptor (aGPCR) GPR126/ADGRG6 plays an important role in several physiological functions, such as myelination or peripheral nerve repair. This renders the receptor an attractive pharmacological target. GPR126 is a mechano-sensor that translates binding of extracellular matrix (ECM) molecules to its N terminus into a metabotropic intracellular signal. To date, the structural requirements and the character of the forces needed for this ECM-mediated receptor activation are largely unknown.

In this study we provide this information by combining classic second messenger detection with single cell atomic force microscopy. We establish a monoclonal antibody targeting the N terminus to stimulate GPR126 and compare it to the activation through its known ECM ligands collagen IV and laminin 211. As each ligand uses a distinct mode of action, the N terminus can be viewed as an allosteric module that can fine-tune receptor activation in a context-specific manner.

## Introduction

The adhesion G protein-coupled receptor (aGPCR) GPR126/ADGRG6 plays an essential role in several important physiologic and pathogenic processes, including myelination (Monk et al., 2009; Monk et al., 2011; Mogha et al., 2013; Ravenscroft et al., 2015), peripheral nerve injury and repair (Mogha et al., 2016), the development of the peripheral nervous system (PNS) (Ravenscroft et al., 2015), and the differentiation of osteoblasts (Sun et al., 2020) as well as adipocytes (Suchý et al., 2020). Further, an association of *GPR126* variants with the development of scoliosis was found in humans and mice (Kou et al., 2013; Karner et al., 2015; Xu et al., 2015; Qin et al., 2017; Kou et al., 2018; Liu et al., 2018; Man et al., 2019; Takeda et al., 2019; Xu et al., 2019c; Xu et al., 2019b; Xu et al., 2019a). Thus, pharmacological targeting of this receptor is of high interest. The physiological implications are mainly attributed to the modulation of cAMP levels by the receptor, which is achieved through the receptor’s coupling to G_s_ protein (Mogha et al., 2013).

Like other aGPCRs GPR126 harbors an endogenous tethered agonistic sequence located distal of the GPS cleavage motif, termed the *Stachel* sequence (Liebscher et al., 2014). Synthetic peptides derived from this integral agonist can be used as agonists on the receptor (Liebscher et al., 2014; Demberg et al., 2017). More recently, small molecule agonists have been identified (Bradley et al., 2019; Diamantopoulou et al., 2019), but similar to the agonistic peptides, they lack specificity for the receptor (Demberg et al., 2017; Bradley et al., 2019). Additional activation can be achieved through the receptor’s N-terminal ligands collagen IV (Paavola et al., 2014), prion protein PrP^C^ (Küffer et al., 2016) and laminin 211 (Petersen et al., 2015). Yet, none of these agonists are specific for GPR126 and ECM proteins lack characteristics of a classic receptor agonist as they are long-lived, stable molecules with essentially no diffusivity (Bassilana et al., 2019; Baxendale et al., 2021). Thus, it is unclear how the interaction between aGPCR and the ECM can be interpreted as a specific signal to modulate receptor activity levels. Mechanical forces are suggested to facilitate this interaction (Liebscher et al., 2014; Stoveken et al., 2015); (Scholz et al., 2017; Dannhäuser et al., 2020), however, the force input as well as the structural components needed for activation have not been defined.

Targeting the large N terminus of an aGPCR with an antibody provides a specific way for receptor activation, which has been successfully shown for two other representatives of this receptor class (Yona et al., 2008; Bhudia et al., 2020; Chatterjee et al., 2021). The mechanism behind this N-terminally mediated signal is currently as unclear as the signals mediated by the ECM ligands. Understanding these fundamental activation processes will not only increase our understanding of the physiological circumstances that govern GPR126-mediated functions but it will set the stage for allosteric pharmaceutical targeting of this and potentially other aGPCRs.

In the absence of a specific antibody recognizing the N terminus of GPR126, we used an antibody recognizing an N-terminal HA epitope in GPR126, which was sufficient to activate the receptor. Our study characterizes the structure-function prerequisites for this antibody-mediated activation and describes in real time the type and strength of mechanical input needed to activate GPR126 through either antibody or endogenous ligands collagen IV and laminin 211. We show that the activation through the antibody is mediated through cross-linking of the receptor, while collagen IV and laminin 211 need specific pushing and pulling forces, respectively. As the occurrence of ECM molecules is timely and locally regulated in tissues a temporo-spatial and force-specific activation of GPR126 can be achieved through the N terminus as allosteric force integrator.

## Results

### An antibody targeting the N-terminal HA epitope can activate GPR126

Since GPR126 can be activated through interaction with its extracellular ligands and mechanical stimuli, it can be assumed that the N terminus plays a decisive role in mediating these signals. However, our understanding of these dynamic processes is limited. In order to establish a specific N-terminal interacting partner of GPR126, we established an antibody-based approach. Due to the lack of antibodies targeting the endogenous GPR126 sequence, we used the hemagglutinin (HA) epitope for our experimental setup and inserted it right after the predicted signal peptide of the receptor (Fig. 1A). Surprisingly, increasing concentrations of the commercially available anti-HA antibody significantly elevated cAMP levels in COS-7 cells transfected with the HA-tagged wild type (WT) GPR126 (Fig. 1B), but not in the empty vector transfected control cells. In a control experiment with an anti-FLAG antibody targeting the C-terminal epitope of GPR126 no change in cAMP levels was observed (Supp. Fig. S1A).

**Figure 1.**
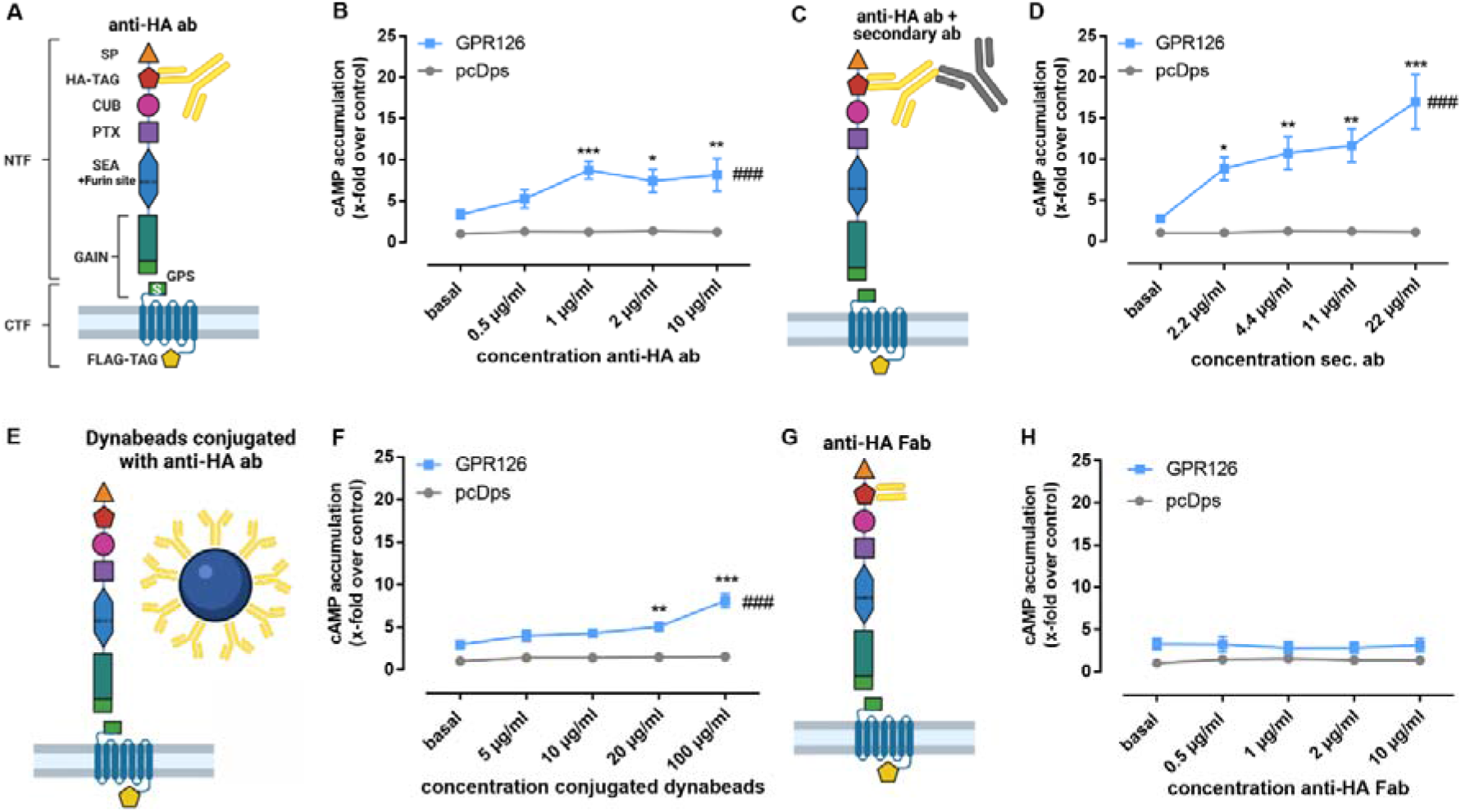
Antibodies against the N terminus activate GPR126. (**A**) Schematic setup for the anti-HA antibody (anti-HA ab) stimulation of full-length WT GPR126 in cAMP accumulation assays. The N terminus contains the signal peptide (SP, orange triangle), a complement C1r/C1s-Uegf-BMP1 domain (CUB, magenta oval), a pentraxin domain (PTX, purple square) and the Sperm protein, Enterokinase and Agrin (SEA) domain (blue hexagon) including the furin site. The highly conserved GAIN domain (green rectangle) contains the GPS, which is followed by the *Stachel* sequence (S). In our receptor constructs, we inserted an N-terminal hemagglutinin tag (HA-TAG, red pentagon) immediately distal to the signal peptide and a C-terminal Flag tag (FLAG-TAG, yellow pentagon) right before the Stop codon. (**B**). Different concentrations of anti-HA antibody (0.5 μg/ml ≙ 3.3 μM; 1 μg/ml ≙ 6.67 μM; 2 μg/ml ≙ 13.34 μM; 10 μg/ml ≙ 66.7 μM) were used to treat vector control (pcDps) and GPR126-transfected COS-7 cells (effect of construct p = 0.0410, effect of concentration p < 0.0001, interaction construct × concentration p = 0.0506; two-way ANOVA). Basal cAMP level in pcDps transfected cells: 8.9 ± 0.8 nM/well. (**C**, **D**) Amplification of cAMP signal of GPR126 with 1 μg/ml of anti-HA ab and subsequent incubation with different concentrations of secondary ab (2.2 μg/ml ≙ 14.7 μM; 4.4 μg/ml ≙ 29.3 μM; 11 μg/ml ≙ 73.3 μM; 22 μg/ml ≙ 146.7 μM) on vector control and GPR126-transfected COS-7 cells (effect of construct p = 0.0104, effect of concentration p < 0.0001, interaction construct × concentration p = 0.0072; two-way ANOVA). Basal cAMP level in pcDps transfected cells: 4.2 ± 1.0 nM/well. (**E**, **F**) Paramagnetic Dynabeads® were conjugated with anti-HA ab and used in different concentrations for stimulation of vector control and GPR126-transfected COS-7 cells (effect of construct p = 0.0001, effect of concentration p < 0.0001, interaction construct × concentration p < 0.0001; two-way ANOVA). Empty vector (pcDps) served as negative control. Basal cAMP level in pcDps transfected cells: 5.8 ± 0.4 nM/well. (**G**, **H**) cAMP accumulation upon incubation with indicated concentrations of Fab fragment (0.5 μg/ml ≙ 10 μM; 1 μg/ml ≙ 20 μM; 2 μg/ml ≙ 40 μM; 10 μg/ml ≙ 200 μM) on vector control and GPR126-transfected COS-7 cells (effect of construct p < 0.0001, effect of concentration p = 0.9920, interaction construct x concentration p = 0.9032; two-way ANOVA). Basal cAMP level in pcDps transfected cells: 7.7 ± 1.2 nM/well. All data are given as means ± SEM of three – five independent experiments each performed in triplicates. Statistics were performed by applying a two-way ANOVA followed by Dunnett post hoc analysis; *p < 0.05; **p < 0.01; ***p < 0.001. All significances given as stars (*) above individual points in the graphs show the result of the post hoc analysis, while # indicates significant concentration-dependent effects (###p < 0.001). Schematic images were created with BioRender.com.

Adding increasing concentrations of a secondary antibody to a fixed concentration of 1 μg/ml of the anti-HA antibody yielded an even stronger activation of the receptor (Fig. 1C/D). In order to elucidate whether the observed activation is a consequence of receptor cross-linking through the antibodies or due to the additional weight being attached to the receptor’s N terminus, we added anti-HA antibody-conjugated paramagnetic Dynabeads®, which are decisively larger in size than antibodies (Fig. 1E). We observed a significant increase in cAMP levels (Fig. 1F), which was comparable with anti-HA antibody-mediated activation alone (Fig. 1B) but lower than the combination of primary and secondary antibody (Fig. 1D), indicating that simply adding weight is not the sole key to GPR126 activation. In line with this observation, exposure to a 700 Gs magnetic field placed below the cell layer cannot further enhance conjugated Dynabead-mediated activation (Suppl. Fig. S1B). Crosslinking, however, seems to be a key element to anti-HA-antibody-mediated activation as incubation with the respective monomeric Fab fragment cannot induce cAMP accumulation (Fig. 1G/H).

## Anti-HA antibody-mediated activation depends on the *Stachel* sequence and GPS cleavage

The N terminus of GPR126 includes five structurally different domains (Leon et al., 2020), which can be subject to tissue-specific splicing (Knierim et al., 2019) and serve to interact with the known ligands collagen IV (Paavola et al., 2014), laminin 211 (Petersen et al., 2015) and prion protein (Küffer et al., 2016). To investigate their role in mechano-sensing, we used different N-terminal deletion mutants of GPR126, whose basal activity and expression levels were previously reported (Petersen et al., 2015). The constructs generated were: ΔCUB (lacking 109 aa compared to the WT), ΔPTX (lacking 206 aa) and ΔCUBΔPTX (lacking 315 aa). Additionally, we generated an N-terminally prolonged variant of GPR126 containing an mRuby-tag (236 aa prolongation compared to the WT) (Fig. 2A). Reliable cell surface expression levels of the mutants were demonstrated with ELISA (Suppl. Fig. S2A). The ability of receptor activation for each mutant was measured in cAMP accumulation assays using the synthetic GPR126 *Stachel* peptide as stimulus, thereby ensuring undisturbed signaling (Suppl. Fig. S2B). We stimulated these mutants with the same antibody-based setup as described in Fig. 1 and found that deletion of the CUB and PTX domains yields results highly similar to WT GPR126 (Fig. 2B). The N-terminally prolonged mRuby construct, despite showing WT-like expression and peptide activation (Suppl. Fig. S2A/B), could not be activated through antibodies (Fig. 2B).

**Figure 2.**
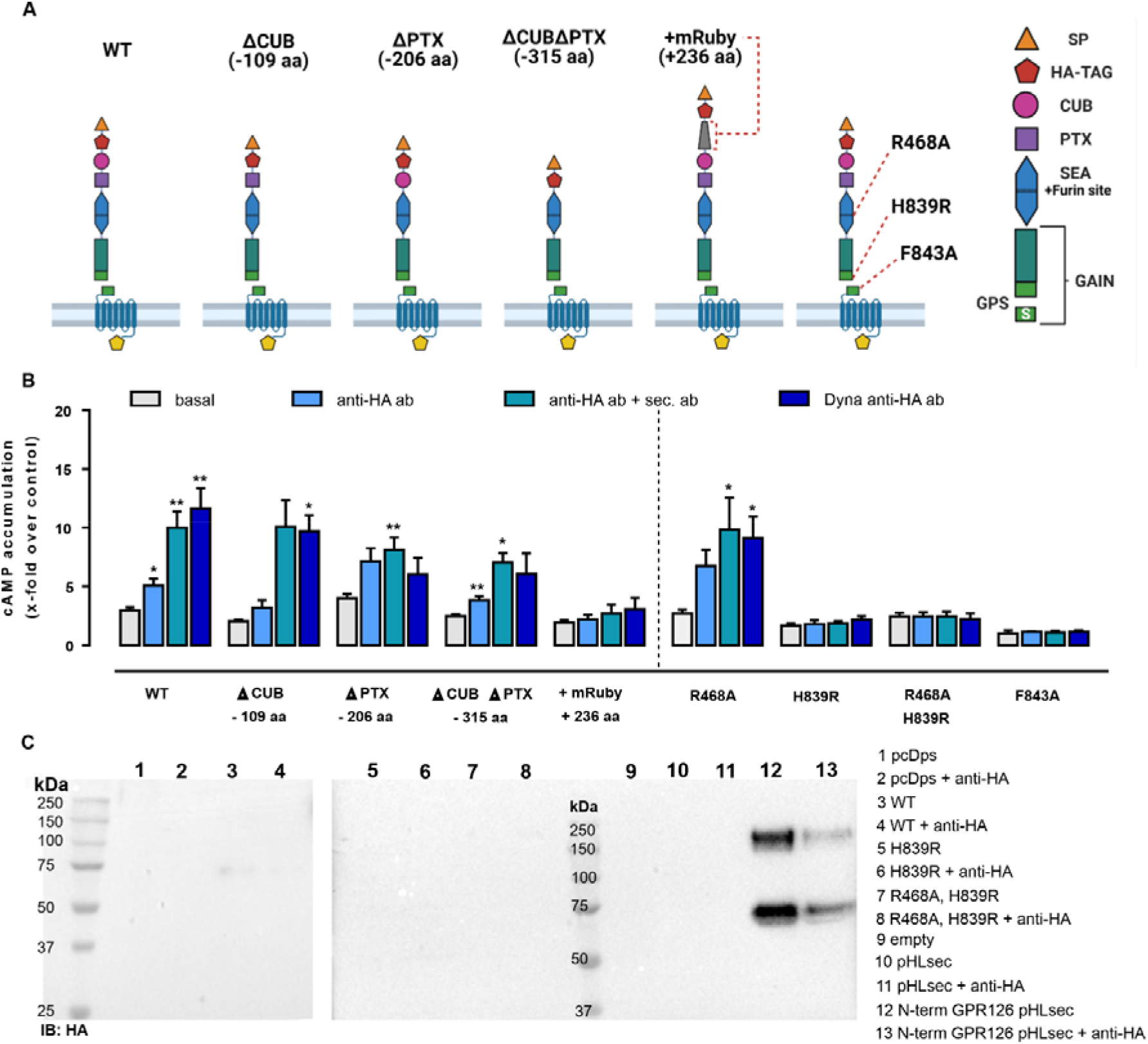
Antibody-mediated activation of GPR126 depends on autoproteolytic cleavage and is obliterated in a construct with an elongated N terminus. (**A**) The domain architecture of the human GPR126 WT and respective mutants is depicted. The positions of the furin deficient receptor mutant R468A, the GPS cleavage mutant H839R and the tethered agonist mutant F843A within the N terminus are displayed. Images were created with BioRender.com. (**B**) The receptor variants were transfected into COS-7 cells and analyzed with the same antibody-based approach as used in Fig. 1 and tested in cAMP assays. Empty vector (pcDps) served as negative control (cAMP level: 4.3 ± 0.7 nM/well). Data are given as mean ± SEM of three – eight different experiments each performed in triplicates. Statistics were performed by applying one-way ANOVA followed by Dunnett post hoc analysis; *p < 0.05; **p < 0.01; ***p < 0.001. All shown statistical significances compare the condition to the basal cAMP accumulation of the same construct. (**C**) COS-7 cells were transfected with the indicated constructs and treated with 1 μg/ml mouse anti-HA ab for 1 h, 48 h after transfection. Then, supernatants were harvested and analyzed by Western blot. Membranes were incubated with rabbit anti-HA ab and secondary HRP conjugated anti-rabbit ab. Corresponding Western blot of cell lysates can be found in Fig. S2C. Secreted N-terminal fragment of GPR126 was only detected when the secretion vector pHLsec containing just the N-terminal fragment of GPR126 as positive control was transfected.

The complex architecture of the N terminus of GPR126 includes two cleavage sites; the GPCR proteolysis site (GPS) within the GAIN domain (Arac et al., 2012) at which the receptor is cleaved into an N-terminal fragment (NTF) and a C-terminal fragment (CTF) at the conserved HLT motif (Lin et al., 2004; Moriguchi et al., 2004), and a furin site, located in the SEA domain (Fig. 2A). To analyze whether cleavage at either position may be required for antibody-mediated activation, we generated the furin-cleavage-deficient receptor mutant R468A, the GPS-cleavage-deficient mutant H839R, and the double-deficient mutant R468A H838R to test in antibody-mediated activation. Proper protein expression, activation capacity through *Stachel* peptide as well as cleavage-deficiency of the mutants was confirmed (Suppl. Fig. S2A-C). While mutation of the furin site (R468A) has no effect on the antibody-mediated stimulation approach, deletion of the GPS cleavage (H839R) abolishes signaling capacity similar to the previously described tethered agonist mutant F843A (Liebscher et al., 2014).

This observation could support the notion that antibody-mediated activation of this aGPCR requires the dissociation of the NTF from the CTF, as has been suggested for the activation of GPR133 (Frenster et al., 2021). To test this assumption, we harvested the supernatants from empty vector and GPR126 transfected cells with our without anti-HA antibody stimulation and subjected them to Western blot analysis (Fig. 2C). As a positive control we used the N terminus of GPR126 encoded on the secretion vector pHLsec. No soluble NTF was found in the supernatants except for the positive control (Fig. 2C). However, a faint band was visible for the WT GPR126 of approximate 75 kDa, suggesting residual removal of the N terminus due to furin cleavage as seen for the cell lysate of the WT construct (Suppl. Fig. S2C). We observed no increase in band intensities after prior stimulation with the mouse anti-HA antibody. Thus, even though the autoproteolytic procession of GPR126 at the GPS site is a prerequisite for antibody-mediated activation it does not lead to NTF removal. It is conceivable that proteolytic procession at the GPS results in a favorable orientation of the *Stachel* sequence, which is indispensable for GPR126 activity.

### Anti HA antibody activation of GPR126 does not require additional pushing and pulling forces

To evaluate whether cross-linking alone is sufficient or if additional mechanical forces are needed to activate GPR126 through the anti-HA antibody, we employed an atomic force microscopy (AFM) approach. We theorized that the antibody could exert either pulling (through cross-linking) or pushing forces (through residing on the cell layer) on the receptor, and that both can be quantified with AFM. To do so, tipless AFM cantilevers were coated with anti-HA antibody using PEG linkers according to a well-established protocol (Ebner et al., 2007). In contrast to cantilevers with a tip, this approach allows for multiple antibodies to bind and thus interact with the cell surface. Therefore, the mechanical force induced by the cantilever and mediated by the antibodies is applied to multiple receptors expressed on the cell surface simultaneously. The coated cantilevers were then used to apply pressure (pushing) or a force-clamp (pulling) to individual cells that were successfully co-transfected with the given receptor construct and the Pink Flamindo cAMP sensor (Harada et al., 2017) (Fig. 3A). This setup allows for the simultaneous application of force and the detection of changes in intracellular cAMP levels. Single cells within a confluent monolayer were chosen for the measurements based on the detection of the GFP signal from the pUltra vector, which allows for bicistronic expression of EGFP and GPR126 (Lou et al., 2012) and the fluorescent signal from the Pink Flamindo cAMP sensor (Harada et al., 2017) (Suppl. Fig. S3A). Since COS-7 cells are not suited for AFM experiments due to their weak adherence to glass coverslips, we used GripTite™ 293 MSR (HEK-GT) cells instead. Receptor cell surface expression and activation in HEK-GT cells was confirmed prior to AFM experiments by ELISA and cAMP accumulation assays, respectively (Suppl. Fig. S3B/C). Relative possible Pink Flamindo fluorescence changes upon stimulation of GPR126 with either forskolin, anti-HA antibody or *Stachel* peptide pGPR126 are displayed in Suppl. Fig. S3D.

**Figure 3.**
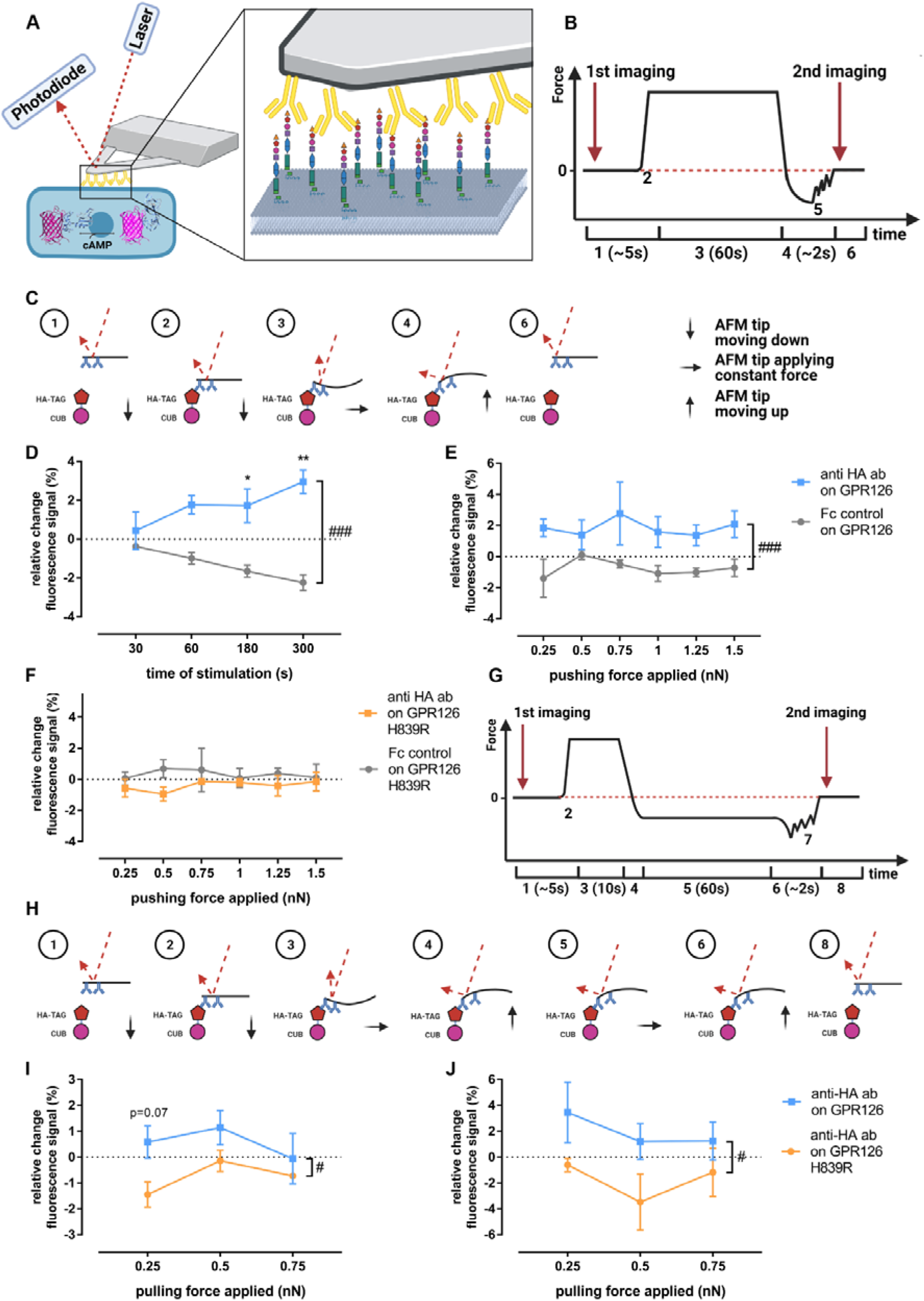
Evaluating potential mechano-activation of GPR126 via anti-HA antibodies. (**A**) Schematic AFM experiment setup with a coated tipless cantilever pressing on a cell co-transfected with GPR126 and the Pink Flamindo cAMP sensor. (**B, C)** Course of the AFM cantilever deflection during the pushing experiments and the forces applied to the receptor and corresponding laser deflection: (1) cantilever approaches cell; (2) point of contact between cantilever and cell; (3) constant pressure being applied to the cell; (4) cantilever is retracted from the cell; (5) rupture of cantilever bindings; (6) all bindings are ruptured and the cantilever is back in its starting position. (**D**) Changes in Pink Flamindo fluorescence intensity after applying a pushing force of 1 nN on the WT receptor at different time points (effect of time p = 0.775, effect of coating p = 0.0002, interaction time × coating p = 0.065; two-way ANOVA). (**E, F**) Changes in Pink Flamindo fluorescence intensity after applying indicated pushing forces for 60s on the WT receptor (**E**) (effect of force applied p = 0.878, effect of coating p < 0.001, interaction force × coating p = 0.882; two-way ANOVA) and the H839R mutant (**F**) (effect of force applied p = 0.981, effect of coating p = 0.0867, interaction force × coating p = 0.924; two-way ANOVA). (**G, H**) Course of the AFM cantilever deflection during the force clamp experiments and the forces applied to the receptor and corresponding laser deflection: (1) cantilever approaches cell; (2) point of contact between cantilever and cell; (3) initial pressure being applied to allow binding between cantilever and cell; (4) cantilever is retracted from the cell until the desired pulling force is applied; (5) constant pulling force is applied; (6) cantilever is retracted from the cell completely; (7) rupture of cantilever bindings; (8) all bindings are ruptured and the cantilever is back in its starting position. Changes in Pink Flamindo fluorescence intensity after applying indicated pulling forces over (**I**) 60s (effect of force applied p = 0.297, effect of receptor variant p = 0.0135, interaction force × receptor variant p = 0.497; two-way ANOVA) and (**J**) 300s (effect of force applied p = 0.405, effect of receptor variant p = 0.0386, interaction force × receptor variant p = 0.804; two-way ANOVA). Data are given as means ± SEM of three-five different experiments each measuring three individual cells for each force and time. Statistics were performed as two-way ANOVA followed by Tukey post hoc analysis; *p < 0.05; **p<0.01. All significances given as stars (*) above individual points in the graphs show the result of the post hoc analysis, while # indicates significant coating-dependent (D and E) or receptor-variant-dependent (I and J) effects (#p < 0.05; ###p<0.001). Schematic images (A-C and G-H) were created with BioRender.com. The Pink Flamindo depiction in A was taken from (Harada et al., 2017).

The course of the cantilever deflection and therefore the force applied to the cell for the pushing setup is depicted in Figures 3B and 3C. The cAMP-evoked changes in the Pink Flamindo fluorescence were monitored by imaging right before the AFM cantilever applied pressure or a pulling force and immediately after its retraction (Fig. 3B). A cantilever coated with human Fc-protein instead of anti-HA antibodies was used as negative control.

When a pushing force of 1 nN was applied over varying times, a significant increase in the Pink Flamindo fluorescence signal could be observed in GPR126-transfected cells compared to the negative control. This effect got stronger and more significant over time suggesting continuous stimulation of the receptor (Fig. 3D). The control condition responded with a reduction of cAMP sensor intensity over time, presumably due to bleaching effects induced by the AFM detection laser. Having established that a significant increase in intracellular cAMP can be achieved by pressure application via anti-HA antibody coated tips, we applied varying forces over a constant time in order to quantify the strength of the pushing force needed to activate GPR126 (Fig. 3E). Applying varying pushing forces (0.25nN - 1.5nN) with anti-HA antibody coated tips over 60s lead to a significant increase in cAMP levels consistently over the whole range of applied forces. This indicates that either the binding, respective cross-linking, of the antibody to the receptor or the pushing forces that occur during the encounter between coated cantilever and receptor are already sufficient to activate GPR126. When we performed the same experiment with the cleavage-deficient mutant H839R (Fig. 3F), which showed no activation in cAMP accumulation assays using antibodies and dynabeads (Fig. 2B), we again observed no change in the cAMP-mediated Pink Flamindo signal regardless of the pressure applied.

To investigate how pulling forces affect GPR126 activation, we used the force-clamp setup as shown in Figs. 3G and H. An Fc-control antibody-coated cantilever could not be used as a negative control in this case, as it does not bind to the receptor or the HA-epitope and therefore no pulling forces would be applied. Instead we compared the WT receptor to the insensitive H839R mutant, which was still able to bind the anti-HA antibody. We applied varying pulling forces (0.25 nN - 0.75 nN) over 60s (Fig. 3I) and 300s, as fold changes were very low after the shorter pulling time period (Fig. 3J). There was a significant difference in the responses observed in WT and cleavage deficient receptor, however, highest fluorescence signals were detected for the lowest pulling force (0.25 nN), while an increase in this stimulus tended to reduced cAMP production. The magnitude of the observed Pink Flamindo signal corresponds to those seen in the pushing approach. Thus, the detected increase under low pulling conditions might be due to the same reasons as the observed pushing signal (cross-linking or initial interaction push) while applying stronger pulling forces might actually inactivate the receptor.

### Endogenous ligands of GPR126 convey a highly specific type of mechano-activation

We established a reliable way to apply defined mechanical pulling and pushing forces on GPR126 and measured the following relative changes in intracellular cAMP levels. We set to investigate the forces needed to stimulate GPR126 using its natural ligands. Several ligands have been shown to modulate GPR126 activity (Fig. 4A): collagen IV has been described as directly activating ligand (Paavola et al., 2014), which could be interpreted as a pushing force on the receptor as it ‘sits’ on it. Laminin 211 on the other hand was reported to require mechanical stimuli such as shaking or vibration to induce cAMP signaling (Petersen et al., 2015), which could be a proxy for pulling forces. To test these assumptions, tipless AFM cantilevers were coated with collagen IV or laminin 211.

**Figure 4.**
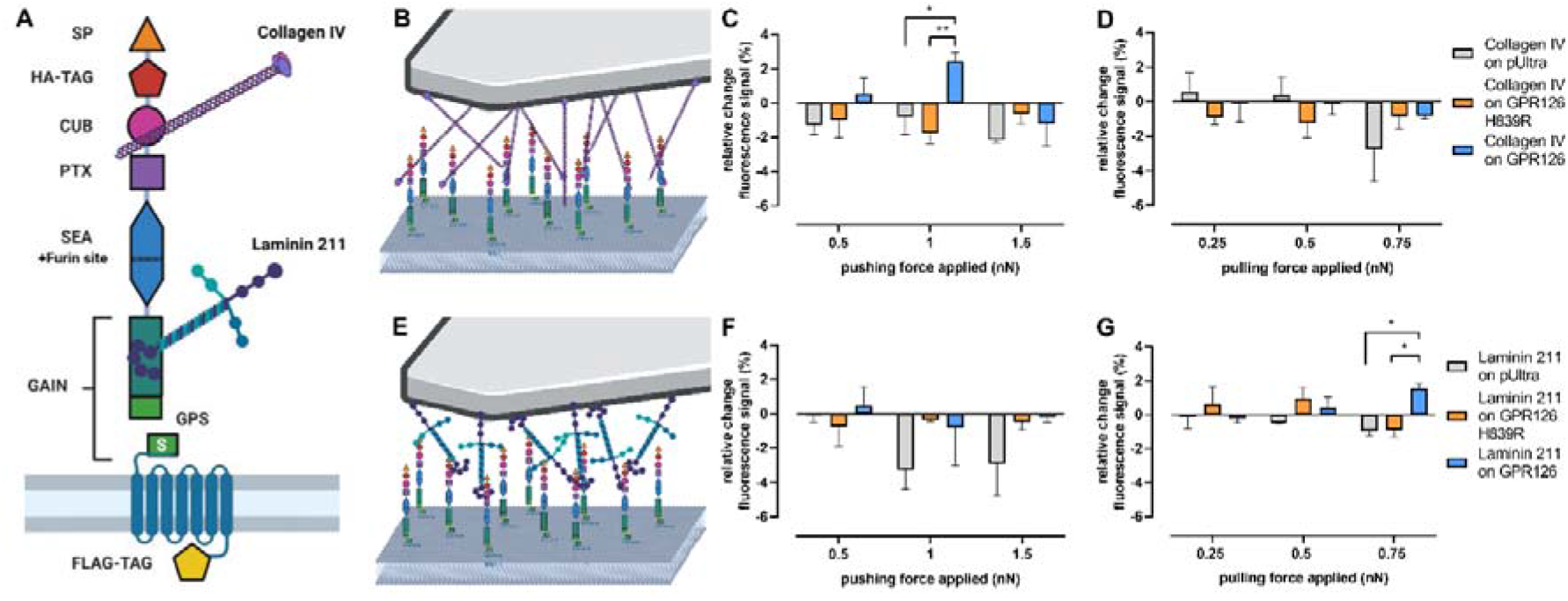
Mechano-activation of GPR126 via its ligands. (**A**) The domain architecture of human GPR126 with the binding sites of its ligands, collagen IV and laminin 211, is depicted. (**B, E**) Schematic demonstrating the AFM setup with (**B**) collagen IV- and (**E**) laminin 211-coated cantilevers. (**C, D**) Changes in Pink Flamindo fluorescence intensity after applying a varying pushing (**C**) or pulling (**D**) force over 60s with a collagen IV-coated cantilever. (**F, G**) Changes in Pink Flamindo fluorescence intensity after applying a pushing (**F**) or pulling (**G**) force over 60s with a laminin 211-coated cantilever. Data are given as means ± SEM of three different experiments, each measuring three individual cells for each force. Statistics were performed as two-way ANOVA (**C**: effect of construct p = 0.1452, effect of force applied p = 0.0635, interaction construct × force applied p = 0.0298; **D**: effect of construct p = 0.8096, effect of force applied p = 0.1748, interaction construct × force applied p = 0.2744; **F**: effect of construct p = 0.0537, effect of force applied p = 0.4536, interaction construct × force applied p = 0,7286; **G**: effect of construct p = 0.2225, effect of force applied p = 0.5884, interaction construct × force applied p = 0.0246) in combination with Tukey post hoc analysis; *p < 0.05; **p < 0.01. Significances in the graphs show the results of the post hoc analysis. Schematic images were created with BioRender.com

When pushing at WT GPR126 transfected cells with a collagen IV-coated cantilever (Fig. 4B), it took specifically 1 nN to induce a significant increase in cAMP levels (Fig. 4C). Lower or higher pressure did not activate the receptor. Again, GPR126 H839R could not be activated through this mechanical stimulus. Applying pulling forces with collagen IV did not produce any significant changes in the Pink Flamindo fluorescence signal for neither the WT receptor nor the cleavage-deficient mutant (Fig. 4D). The laminin 211-coated cantilever (Fig. 4E) did not activate GPR126 through pushing (Fig. 4F), but a significant increase in the Pink Flamindo fluorescence signal was seen in the force-clamp setup when applying a pulling force of 0.75 nN (Fig. 4G). Lower pulling forces were not sufficient to activate the receptor.

## Discussion

The aGPCR GPR126 can be activated through different mechanisms, including agonistic *Stachel* sequence-derived peptides (Liebscher et al., 2014), its ligands collagen IV (Paavola et al., 2014), laminin 211 (Petersen et al., 2015) and prion protein (Küffer et al., 2016), the small molecule compound apomorphine (Bradley et al., 2019) and mechanical stimuli such as vibration or shaking (Petersen et al., 2015). Yet, all of these activators lack specificity for GPR126, which hampers their practicality as tools for *in vivo* experiments or potential therapeutic approaches. The agonistic peptide pGPR126 can for example cross-activate the aGPCR GPR64/ADGRG2 (Demberg et al., 2017), while the ECM molecules collagen and laminin can also bind and activate integrins (Keely et al., 1995) and apomorphin is an agonist on dopamine receptors (Millan et al., 2002). In this study, we show that a commonly used monoclonal antibody targeting an N-terminal HA epitope can serve as an activator of GPR126.

Antibody-mediated GPCR activation is a known phenomenon. For example, in several disease contexts, autoantibodies target GPCRs, such as the thyroid stimulating hormone receptor, calcium-sensing receptor and muscarinic M1 and M2 receptors (Unal et al., 2012). Previously, the aGPCRs EMR2 and GPR56 were shown to be activated through antibodies targeting the receptors’ N-termini (Yona et al., 2008; Bhudia et al., 2020; Chatterjee et al., 2021). In the absence of known antibodies against GPR126, we probed a commercial anti-HA antibody targeting an artificially introduced HA epitope at the N terminus of GPR126 and found that this universal antibody was indeed capable of activating the receptor (Fig. 1B). We, thus, wondered how the anti-HA antibody mediates this activation. Agonistic properties of GPCR-targeting antibodies have been previously assigned to their interaction with the cognate ligand’s binding pocket or stabilization of ligand-induced active receptor conformations (Gupta et al., 2008) for example through cross-linking/dimerization of the receptor as has been described for β1-adrenergic (β_1_AR) receptor (Hutchings et al., 2014). With respect to the known and anticipated activation scenarios for aGPCRs, it would also be conceivable that the antibody leads to a dissociation of the NTF resulting in exposure of the tethered agonist (Petersen et al., 2015; Mathiasen et al., 2020; Frenster et al., 2021) or that it mediates mechanical stimuli such as pushing or pulling.

Our results support the notion that the anti-HA antibody-mediated activation is most likely due to cross-linking of the receptor as a monomeric Fab fragment of the anti-HA antibody alone is not able to activate GPR126 (Fig. 1H). The observation that addition of a secondary antibody further enhanced cAMP production (Fig. 1D) indicated that the receptor might also respond to the weight of molecules pushing onto it. However, neither paramagnetic Dynabeads® coated with the anti-HA antibody alone (Fig. 1F) nor in combination with a magnet below the cell monolayer (suppl. Fig. 1B) enhanced cAMP levels compared to anti-HA antibody incubation alone. These results as well as a lack of activation through direct pushing or pulling with an anti-HA antibody-coated cantilever in the AFM setup (Fig. 3) demonstrate that cross-linking through the antibody is sufficient to activate GPR126 while no additional forces are required. Our mutagenesis data shows that the anti-HA antibody-mediated stimulation of GPR126 depends on an intact tethered agonist sequence, thus we can rule out the option that the antibody can interact with the endogenous agonist binding pocket (Fig. 2B). Similarly, cleavage at the GPS is essential for this activation (Fig. 2B), but we found no indication that this would lead to a dissociation of the NTF (Fig. 2C). It can be speculated that this cleavage event would be required instead to induce a conformation that is necessary for the tethered agonist to reach its binding pocket. This is in contrast to other aGPCRs like GPR56 (Chatterjee et al., 2021) and latrophilin (Scholz et al., 2015), whose antibody- and mechano-mediated activations, respectively, are not affected by mutations of their GPS cleavage motifs. Thus, it seems that autoproteolysis at the GPS serves distinct purposes among different receptors.

When studying aGPCR activation by ligands or antibodies, splice variants normally have to be considered, since the complex exon-intron composition results in a large subset of functionally divergent receptors. As an example, activation of GPR56 through an antibody targeting the GAIN domain is dependent on a Serine–Threonine–Proline-rich (STP) region, which otherwise does not influence basal signaling levels of the receptor (Chatterjee et al., 2021). In the case of GPR126, alternative splicing influences the domain composition of the N terminus (Knierim et al., 2019). Within the N terminus, the CUB/PTX domain is of functional relevance as it serves as point of interaction for collagen IV (Paavola et al., 2014), yet, for anti-HA antibody-mediated activation it appears to be neglectable. In contrast, elongating the receptor’s N terminus through the addition of a fluorescent protein abolishes this activation. Similar observations have been made for the adhesion GPCR dCIRL, where elongation of the N terminus reduces the response to mechanical stimuli (Scholz et al., 2017). Thus, large artificial N-terminal domains influence signaling properties, even when they do not act as binding sites for the activating antibodies. It is currently unknown whether this is due to the simple change in length or an altered three-dimensional structure of the N terminus.

The endogenous interaction partners collagen IV and laminin 211 also bind to the N terminus of GPR126 but show different activation mechanisms. While incubation with collagen IV directly activates the receptor (Paavola et al., 2014), laminin 211 only induces cAMP production in combination with mechanical forces (Petersen et al., 2015). As the quality and the quantity of the required forces have not been defined on a single cell level it was hard to judge whether the *in vitro* findings could possibly be relevant in an *in vivo* context. Using the AFM approach we found that laminin 211 requires increasing pulling forces of at least 0.75 nN (Fig. 4G), while collagen IV only raises cAMP levels upon a pushing force of 1 nN (Fig. 4C). Both mechano-stimulations require a cleavable GPR126. Both activation patterns fit the physiological setting for these ligands in the process of myelination (Bunge et al., 1990; Paavola et al., 2014; Petersen et al., 2015) and the detected forces needed to induce a ligand-specific response are within the physiologic force range. They are below the traction forces that are normally transmitted by cell adhesions to the surrounding ECM (1 – 10 nN) (Balaban et al., 2001), indicating that already small changes can be detected by an aGPCR.

In summary, we were able to define the different ways in which a subset of structurally divergent molecules binding to the N terminus of GPR126 can modify the activity of this receptor. This establishes the N terminus as an allosteric integrator of signals coming from the immediate extracellular surrounding that is able to induce a spatio-temporal-force-dependent signal. It should therefore be considered as prime target for future pharmaceutical interventions as it provides the basis for receptor and potentially signalling-specific modulation of GPR126 activity.

## Materials and Methods

If not stated otherwise, all standard substances were purchased from Sigma Aldrich (Taufkirchen, Germany), Merck (Darmstadt, Germany), and C. Roth GmbH (Karlsruhe, Germany). Cell culture material was obtained from Thermo Fisher Scientific (Schwerte, Germany) and primers were obtained from Microsynth Seqlab GmbH (Göttingen, Germany). The magnetic stimulator was created via embedding N45 NdFeB magnets (https://www.supermagnete.de/data_sheet_S-08-05-N.pdf) into a custom 3D printed holder matching the outer diameter of a corresponding cell culture plate.

### Plasmid generation

The constructs of human GPR126 and ΔCUB, ΔPTX, ΔCUBΔPTX mutants have been described previously (Mogha et al., 2013; Liebscher et al., 2014; Petersen et al., 2015). Point mutations for R468A and H839R constructs were inserted by quick change mutagenesis. In brief, plasmid DNA was amplified with PCR and then digested with DpnI restriction enzyme for 4 h at 37°C prior heat shock transformation in *E. coli*. The mRuby epitope was inserted in the human GPR126 construct after the hemagglutinin (HA) epitope by a PCR-based site-directed mutagenesis and fragment replacement strategy. The sequences of all generated mutants of human GPR126 were verified by Sanger sequencing (Microsynth Seqlab, Göttingen, Germany). The GPR126/pULTRA construct was generated using One-step isothermal DNA assembly (Gibson et al., 2009). The reaction buffer was prepared according to protocol. All enzymes were acquired from NEB: T5 exonuclease (M0363), Taq DNA Ligase (M0208) and Phusion DNA Polymerase (M0530). The WT receptor DNA for the assembly was obtained via PCR using the previously described construct of the full-length human GPR126 in the pcDps vector (Liebscher et al., 2014). The pULTRA (pUltra was a gift from Malcolm Moore, Addgene plasmid #24129, (Lou et al., 2012)) vector was restriction digested using Xba1 (NEB, R0145).

### Anti-HA Fab fragment generation

Sequences encoding Anti-HA Fab (clone 12CA5) heavy and light chains were codon optimized for human cells, synthesized and cloned into pcDNA3.4 (Thermo Fisher Scientific) by Genscript. DNA of heavy and light chains was mixed 1:1, transfected into Expi293 cells (Thermo Fisher Scientific) using PEI Max (Polysciences) and expressed at 37°C for 7 days. Fab was purified from clarified culture supernatants using a CaptureSelect CH1-XL column (Thermo Fisher Scientific) and buffer-exchanged into Tris-buffered saline (100 mM NaCl, 20 mM Tris, pH7.5) using a HiPrep 26/10 desalting column (Cytiva, Freiburg, Germany). Binding of Fab fragment to the present HA-tag was proven concentration-dependently on P2Y_12_-transfected cells, which served as receptor expression positive control (Suppl. Fig. S1C).

### cAMP accumulation assays

The human N- and C-terminally tagged GPR126 was heterologously expressed in COS-7 cells grown in Dulbecco’s minimum essential medium (DMEM) supplemented with 10% fetal bovine serum (FBS), 100 units/ml penicillin, and 100 μg/ml streptomycin or the GripTite 293 MSR Cell Line (Thermo Fisher Scientific) (HEK-GT) grown in DMEM supplemented with 10% fetal bovine serum, 1% G418 (Thermo Fisher Scientific, 10131035) and 1% Non-Essential Amino Acids (Thermo Fisher Scientific, 11140050) at 37°C and 5% CO_2_ in a humidified atmosphere. Cells were split into 48-well plates (3×10^4^ cells/well for COS-7 or 1.3×10^5^ cells/well for HEK-GT) for antibody stimulation assays or into 96-well plates (1.5×10^4^ cells/well for COS-7 or 4.5×10^4^/well for HEK-GT) for peptide stimulation. Transfection was done with Lipofectamine 2000 (Thermo Fisher Scientific) according to the manufacturer’s protocol using 50 ng (96-well plates) or 500 ng (48-well plates) of receptor plasmid DNA/well. 48 h after transfection, GPR126 and empty vector-transfected cells were stimulated with the indicated concentrations of anti-HA antibody (H3663, stock concentration 1 mg/ml, Sigma-Aldrich), anti-HA Fab fragment (stock concentration 1 mg/ml, kind gift from T. Schiffner), anti-HA conjugated super paramagnetic Dynabeads® (14311D, stock concentration 10 mg/ml, Thermo Fisher Scientific) or anti-FLAG antibody (F1804, Sigma-Aldrich) in DMEM for 1 h, followed by incubation with 3-isobutylmethyl-xanthine (1 mM)-containing medium or including the secondary antibody anti-mouse IgG (Fc specific, M2650, Sigma-Aldrich) for 30 min. For peptide stimulation, *Stachel*-sequence derived peptide was diluted in IBMX-containing medium. Peptide solution from purified powder was achieved by preparing a 100 mM in 100% DMSO solution, which was further diluted into 10 mM stocks using a 50 mM, pH 8 Tris buffer and finally pH controlled. Peptide concentrations used in assays are 1 mM. The 1 mM peptide solution contains 1% DMSO and 10% of Tris buffer used for dilution. After stimulation cells were lysed in LI buffer (PerkinElmer, Rodgau, Germany) and kept frozen at −80°C until measurement. To measure cAMP concentration, the Alpha Screen cAMP assay kit (PerkinElmer) was used according to the manufacturer’s protocol. The accumulated cAMP was measured in 384-well white OptiPlate microplates (PerkinElmer) with the EnVision Multilabel Reader (PerkinElmer). Super paramagnetic Dynabeads® were conjugated with anti-HA antibody according to the manufacturer’s protocol. Briefly, Dynabeads were weighed, washed with C1 solution, mixed with appropriate amount of anti-HA antibody in C1 solution, C2 solution was added and the mixture was incubated overnight at 37°C on a roller followed by washing steps afterwards.

### Enzyme-linked immunosorbent assay (ELISA)

Cells were split 48-well plates (3×10^4^ cells/well for COS-7 or 1.3×10^5^ cells/well for HEK-GT). To estimate cell surface expression of receptors carrying an N-terminal HA tag an indirect cellular enzyme-linked immunosorbent assay (ELISA) was used (Schöneberg et al., 1998). Briefly, cells were transfected with indicated constructs. 48 h after transfection cells were fixed with 4% formaldehyde, washed with PBS, blocked with 10% FBS medium and incubated with anti-HA POD-conjugated antibody followed by *o*-phenylendiamine treatment. Optical densities were measured at a wavelength of 492 nm with the EnVision Multilabel Reader (PerkinElmer).

### Western blot

For Western blot analysis, cells were split into 24-well plates (6×10^4^ cells/well for COS-7) and transfected with either 250 ng of empty pcDps vector, GPR126, GPR126 R468A or GPR126 R468A/H839R using the standard protocols described earlier. Every other transfected construct was incubated with primary anti-HA antibody for an hour as explained before. Supernatants were harvested and cells were lysed with the addition of 300 μl of 2x SDS loading dye (#S3401, Sigma-Aldrich). After freeze thaw cycling the lysates were run on a 10% SDS-PAGE gel, followed by Western blotting. The PVDF membranes were activated using 100% Methanol and transfer was performed for 1 h at 80 V. Membranes were blocked for 1 h with 5% non-fat dry milk in TBST buffer, washed three times with TBST buffer and incubated over night with either primary antibody (rabbit ant-HA, #3724 and anti-GAPDH, #97166 antibody, Cell Signaling). The following day, membranes were again washed with TBST buffer and incubated with secondary anti-rabbit antibody, which has an HRP conjugation (#7074, Cell Signaling) for 1 h at room temperature. Following three washing steps with TBST buffer, SuperSignal West Pico PLUS Chemiluminescent Substrate (Thermo Fisher Scientific) was added to membranes to visualize protein bands using the Bio-Rad Gel Doc Imager.

### AFM

For all Atomic Force Microscopy (AFM) experiments, HEK-GT cells were cultured at the same conditions as for the *in vitro* functional assays. Cells were seeded on 24 mm glass coverslips (coated with Poly-L-Lysin (Sigma-Aldrich, P4707), (1% PLL solution incubated at 37°C for 5 minutes then washed with PBS and dried under a sterile hood) in 6-well plates (1.5×10^6^ cells/well) and co-transfected with the cAMP sensor Pink Flamindo (Addgene plasmid #102356) and either empty vector or the given GPR126 construct in the pULTRA vector on the next day using Lipofectamine 2000 according to the manufacturer’s protocol. The media was changed to the described culture media ~24 h after transfection and AFM measurements took place ~48 h after transfection: the coverslips were transferred into a 35mm cell culture dish and washed with DMEM without phenol red three times and placed into AFM coverslip holders. 750 μl culture media without phenol red was added and the cells were stored at 37°C and 5% CO_2_ until right before the measurements took place.

Tipless silicon nitride AFM cantilevers (NanoWorld, PNP-TR-TL) were coated with monoclonal anti-HA antibodies produced in mouse (H3663, Sigma-Aldrich) or Fc-control for the antibody-based mechano-activation experiments using flexible PEG spacers as described before (Ebner et al., 2007). Recombinant human Fc was expressed by transfected Chinese hamster ovary (CHO) cells, supernatant was collected and purification was done via His-tag using HisLink Protein Purification Resin (V8823, Promega) according to manufacturer’s instructions. Protein purity was confirmed by Western blot analysis. For the ligand-based mechano-activation experiments, tipless silicon nitride AFM cantilevers were washed twice in Chloroform and dried. The cantilevers were then placed into a collagen IV (C6745, Sigma Aldrich) or laminin-211 (LN221, BioLamina, Sundbyberg, Sweden) solution (0.15 mg/ml) and incubated at 4°C overnight. The next day the cantilevers were washed in HBSS twice and stored in HBSS at 4°C until use.

AFM measurements were performed using a Nanowizard IV AFM (Bruker, Billerica, MA) mounted on an IX 83 inverted optical microscope equipped with a 63x PL APO NA 1.42 oil objective (both Olympus Lifes Sciences, Wallisellen, Switzerland) and coupled to a X-Cite Exacte Light source (Excelitas Technologies, Waltham, MA).

Cantilevers were calibrated using the thermal noise method according to (Slattery et al., 2013).

Successfully double-transfected cells were identified by the GFP from the pULTRA vector and the Pink Flamindo fluorescence signal. The cell was then imaged three times in the following order: Brightfield, GFP and Pink Flamindo using a Zyla sCMOS camera (Andor Technology, Belfast, Northern Ireland). The AFM cantilever was then placed centrally on the cell and approached until contacting the cell surface. Proper positioning was verified by a brightfield image and the baseline Pink Flamindo signal was recorded. During stimulation all light sources except the AFM laser were turned off. Immediately after the stimulation was finished, another image of the Pink Flamindo signal was obtained using the same exposure time as before.

For force clamp measurements, the cantilevers were initially pressed onto the cell with a force of 1 nN for 5 s in order to allow antibody/ligands and receptor to bind. The cantilever was then retracted (1 μm/s) until the desired clamp force was reached. This value was kept constant for the indicated times before the cantilever was fully retracted. Extend and retract length was 15 μm and extend and retract speed was 5 μm/s for all experiments.

For analysis, the Pink Flamindo images made before and after the stimulation were compared by measuring the mean intensity of a rectangular area on the stimulated cell using Fiji Image J (Schindelin et al., 2012). To account for variations affecting all cells independently from the stimulation, such as bleaching, a rectangular area was measured on five other cells that were not touched by the cantilever but expressed Pink Flamindo. The average of the change in these five reference cells was subtracted from the measured change in the stimulated cell to isolate the effect the AFM cantilever stimulation has on the receptor-transfected cell (Suppl. Fig. S3A).

To evaluate the mechano-independent changes of the Pink Flamindo fluorescence signal, HEK-GT cells were split into 96-well plates (4.5×10^4^/well) and transfected the following day analog to the cAMP accumulation assays described earlier. Two days after transfection the media was removed from the wells and 40 μl of DMEM without phenol-red were added to each well. Then the GFP signal (transfection control) as well as the Pink Flamindo fluorescence signal were imaged using a Celigo Image Cytometer (Nexcelom Bioscience). Forskolin, pGPR126 and anti-HA antibody diluted in DMEM without phenol-red were added to a final volume of 50 μl/well and following concentrations: 10 μM forskolin, 1 μM anti-HA antibody and 1 mM pGPR126. The cells were imaged again 60 s after addition of the respective stimulus and the intensity of the Pink Flamindo signal beforehand and afterwards was compared.

### Data Analysis

Receptor expression and activation were analyzed using, one- and two-way ANOVA as well as t-test as indicated at each figure legend. p values < 0.05 were considered statistically significant (*p<0.05; **p<0.01; ***p<0.001). All statistical analyses were performed by GraphPad Prism version 6.00 for Windows (GraphPad Software, Inc., La Jolla, USA) or Microsoft Excel 2016 (Microsoft Corporation, Redmond, USA).

## Acknowledgements

Our research is funded by the German Research Foundation (CRC1052 project number 209933838 to IL, CRC1423 project number 421152132 B05 to IL, FOR2149 project number 246212759, P5) to IL and (STE 2843/1-1 project number) to GS. Further funding comes from the European Union (European Social Fund) to IL and the Swiss National Science Foundation (SNF 310030_197764/1) to VS. Further support was provided by the junior research grant by the Medical Faculty, Leipzig University to CW and by the COST association providing an STSM grant within the COST action CA18240 to JM. DGP was supported by NIH/NINDS R01 NS102665, NYSTEM (NY State Stem Cell Science) IIRP C32595GG, NIH/NIBIB R01 EB028774, NYU Grossman School of Medicine, and DFG FOR2149. We thank Torben Schiffner and Clara T. Schoeder for producing the anti-HA Fab fragment.

## Author Contributions

JM, CW, CS, VS, DGP and IL designed the research plan. JF, GS, JM, HK, SB and CW performed the experiments. JF, GS, JM, SB, CW and IL analyzed results. JM, CW and IL wrote the paper with input from all authors. All authors edited and approved of the manuscript.

## Competing interests

The authors declare no competing interest.

## Notes

### Competing Interest Statement

The authors have declared no competing interest.

### Summary of Updates

We have further investigated the mechanism behind antibody-mediated activation of GPR126 and find that cross-linking of receptors is sufficient induce receptor signaling. We further show that removal of the N-terminal fragment is not required for this process.

## References

Arac, D., Boucard, A.A., Bolliger, M.F., Nguyen, J., Soltis, S.M., Südhof, T.C., and Brunger, A.T. (2012). A novel evolutionarily conserved domain of cell-adhesion GPCRs mediates autoproteolysis. EMBO J 31, 1364-1378.

Balaban, N.Q., Schwarz, U.S., Riveline, D., Goichberg, P., Tzur, G., Sabanay, I., Mahalu, D., Safran, S., Bershadsky, A., and Addadi, L., et al. (2001). Force and focal adhesion assembly: a close relationship studied using elastic micropatterned substrates. Nature Cell Biology 3, 466-472.

Bassilana, F., Nash, M., and Ludwig, M.-G. (2019). Adhesion G protein-coupled receptors: opportunities for drug discovery. Nat Rev Drug Discov 18, 869-884.

Baxendale, S., Asad, A., Shahidan, N.O., Wiggin, G.R., and Whitfield, T.T. (2021). The adhesion GPCR Adgrg6 (Gpr126): Insights from the zebrafish model. Genesis (New York, N.Y. : 2000), e23417.

Bhudia, N., Desai, S., King, N., Ancellin, N., Grillot, D., Barnes, A.A., and Dowell, S.J. (2020). G Protein-Coupling of Adhesion GPCRs ADGRE2/EMR2 and ADGRE5/CD97, and Activation of G Protein Signalling by an Anti-EMR2 Antibody. Sci Rep 10, 1004.

Bradley, E.C., Cunningham, R.L., Wilde, C., Morgan, R.K., Klug, E.A., Letcher, S.M., Schöneberg, T., Monk, K.R., Liebscher, I., and Petersen, S.C. (2019). In vivo identification of small molecules mediating Gpr126/Adgrg6 signaling during Schwann cell development. Annals of the New York Academy of Sciences 1456, 44-63.

Bunge, M.B., Clark, M.B., Dean, A.C., Eldridge, C.F., and Bunge, R.P. (1990). Schwann cell function depends upon axonal signals and basal lamina components. Ann N Y Acad Sci 580, 281-287.

Chatterjee, T., Zhang, S., Posey, T.A., Jacob, J., Wu, L., Yu, W., Francisco, L.E., Liu, Q.J., and Carmon, K.S. (2021). Anti-GPR56 monoclonal antibody potentiates GPR56-mediated Src-Fak signaling to modulate cell adhesion. The Journal of biological chemistry 296, 100261.

Dannhäuser, S., Lux, T.J., Hu, C., Selcho, M., Chen, J.T.-C., Ehmann, N., Sachidanandan, D., Stopp, S., Pauls, D., and Pawlak, M., et al. (2020). Antinociceptive modulation by the adhesion GPCR CIRL promotes mechanosensory signal discrimination. eLife 9.

Demberg, L.M., Winkler, J., Wilde, C., Simon, K.-U., Schön, J., Rothemund, S., Schöneberg, T., Prömel, S., and Liebscher, I. (2017). Activation of Adhesion G Protein-coupled Receptors: AGONIST SPECIFICITY OF STACHEL SEQUENCE-DERIVED PEPTIDES. J Biol Chem 292, 4383-4394.

Diamantopoulou, E., Baxendale, S., La Vega León, A. de, Asad, A., Holdsworth, C.J., Abbas, L., Gillet, V.J., Wiggin, G.R., and Whitfield, T.T. (2019). Identification of compounds that rescue otic and myelination defects in the zebrafish adgrg6 (gpr126) mutant. eLife 8.

Ebner, A., Wildling, L., Kamruzzahan, A.S.M., Rankl, C., Wruss, J., Hahn, C.D., Hölzl, M., Zhu, R., Kienberger, F., and Blaas, D., et al. (2007). A new, simple method for linking of antibodies to atomic force microscopy tips. Bioconjugate chemistry 18, 1176-1184.

Frenster, J.D., Stephan, G., Ravn-Boess, N., Bready, D., Wilcox, J., Kieslich, B., Wilde, C., Sträter, N., Wiggin, G.R., and Liebscher, I., et al. (2021). Functional impact of intramolecular cleavage and dissociation of adhesion G protein-coupled receptor GPR133 (ADGRD1) on canonical signaling. J Biol Chem, 100798.

Gibson, D.G., Young, L., Chuang, R.-Y., Venter, J.C., Hutchison, C.A., and Smith, H.O. (2009). Enzymatic assembly of DNA molecules up to several hundred kilobases. Nat Methods 6, 343-345.

Gupta, A., Heimann, A.S., Gomes, I., and Devi, L.A. (2008). Antibodies against G-protein coupled receptors: novel uses in screening and drug development. Combinatorial chemistry & high throughput screening 11, 463-467.

Harada, K., Ito, M., Wang, X., Tanaka, M., Wongso, D., Konno, A., Hirai, H., Hirase, H., Tsuboi, T., and Kitaguchi, T. (2017). Red fluorescent protein-based cAMP indicator applicable to optogenetics and in vivo imaging. Sci Rep 7, 7351.

Hutchings, C.J., Cseke, G., Osborne, G., Woolard, J., Zhukov, A., Koglin, M., Jazayeri, A., Pandya-Pathak, J., Langmead, C.J., and Hill, S.J., et al. (2014). Monoclonal anti-β1-adrenergic receptor antibodies activate G protein signaling in the absence of β-arrestin recruitment. mAbs 6, 246-261.

Karner, C.M., Long, F., Solnica-Krezel, L., Monk, K.R., and Gray, R.S. (2015). Gpr126/Adgrg6 deletion in cartilage models idiopathic scoliosis and pectus excavatum in mice. Hum Mol Genet 24, 4365-4373.

Keely, P.J., Wu, J.E., and Santoro, S.A. (1995). The spatial and temporal expression of the alpha 2 beta 1 integrin and its ligands, collagen I, collagen IV, and laminin, suggest important roles in mouse mammary morphogenesis. Differentiation 59, 1-13.

Knierim, A.B., Röthe, J., Çakir, M.V., Lede, V., Wilde, C., Liebscher, I., Thor, D., and Schöneberg, T. (2019). Genetic basis of functional variability in adhesion G protein-coupled receptors. Scientific reports 9, 11036.

Kou, I., Takahashi, Y., Johnson, T.A., Takahashi, A., Guo, L., Dai, J., Qiu, X., Sharma, S., Takimoto, A., and Ogura, Y., et al. (2013). Genetic variants in GPR126 are associated with adolescent idiopathic scoliosis. Nat Genet 45, 676-679.

Kou, I., Watanabe, K., Takahashi, Y., Momozawa, Y., Khanshour, A., Grauers, A., Zhou, H., Liu, G., Fan, Y.-H., and Takeda, K., et al. (2018). A multi-ethnic meta-analysis confirms the association of rs6570507 with adolescent idiopathic scoliosis. Sci Rep 8, 11575.

Küffer, A., Lakkaraju, A.K.K., Mogha, A., Petersen, S.C., Airich, K., Doucerain, C., Marpakwar, R., Bakirci, P., Senatore, A., and Monnard, A., et al. (2016). The prion protein is an agonistic ligand of the G protein-coupled receptor Adgrg6. Nature 536, 464-468.

Leon, K., Cunningham, R.L., Riback, J.A., Feldman, E., Li, J., Sosnick, T.R., Zhao, M., Monk, K.R., and Araç, D. (2020). Structural basis for adhesion G protein-coupled receptor Gpr126 function. Nat Commun 11, 194.

Liebscher, I., Schön, J., Petersen, S.C., Fischer, L., Auerbach, N., Demberg, L.M., Mogha, A., Cöster, M., Simon, K.-U., and Rothemund, S., et al. (2014). A tethered agonist within the ectodomain activates the adhesion G protein-coupled receptors GPR126 and GPR133. Cell Rep 9, 2018-2026.

Lin, H.-H., Chang, G.-W., Davies, J.Q., Stacey, M., Harris, J., and Gordon, S. (2004). Autocatalytic cleavage of the EMR2 receptor occurs at a conserved G protein-coupled receptor proteolytic site motif. J Biol Chem 279, 31823-31832.

Liu, G., Liu, S., Lin, M., Li, X., Chen, W., Zuo, Y., Liu, J., Niu, Y., Zhao, S., and Long, B., et al. (2018). Genetic polymorphisms of GPR126 are functionally associated with PUMC classifications of adolescent idiopathic scoliosis in a Northern Han population. J Cell Mol Med 22, 1964-1971.

Lou, E., Fujisawa, S., Morozov, A., Barlas, A., Romin, Y., Dogan, Y., Gholami, S., Moreira, A.L., Manova-Todorova, K., and Moore, M.A.S. (2012). Tunneling nanotubes provide a unique conduit for intercellular transfer of cellular contents in human malignant pleural mesothelioma. PLoS ONE 7, e33093.

Man, G.C.-W., Tang, N.L.-S., Chan, T.F., Lam, T.P., Li, J.W., Ng, B.K.-W., Zhu, Z., Qiu, Y., and Cheng, J.C.-Y. (2019). Replication Study for the Association of GWAS-associated Loci With Adolescent Idiopathic Scoliosis Susceptibility and Curve Progression in a Chinese Population. Spine 44, 464-471.

Mathiasen, S., Palmisano, T., Perry, N.A., Stoveken, H.M., Vizurraga, A., McEwen, D.P., Okashah, N., Langenhan, T., Inoue, A., and Lambert, N.A., et al. (2020). G12/13 is activated by acute tethered agonist exposure in the adhesion GPCR ADGRL3. Nature chemical biology 16, 1343-1350.

Millan, M.J., Maiofiss, L., Cussac, D., Audinot, V., Boutin, J.-A., and Newman-Tancredi, A. (2002). Differential actions of antiparkinson agents at multiple classes of monoaminergic receptor. I. A multivariate analysis of the binding profiles of 14 drugs at 21 native and cloned human receptor subtypes. The Journal of pharmacology and experimental therapeutics 303, 791-804.

Mogha, A., Benesh, A.E., Patra, C., Engel, F.B., Schöneberg, T., Liebscher, I., and Monk, K.R. (2013). Gpr126 functions in Schwann cells to control differentiation and myelination via G-protein activation. J Neurosci. 33, 17976-17985.

Mogha, A., Harty, B.L., Carlin, D., Joseph, J., Sanchez, N.E., Suter, U., Piao, X., Cavalli, V., and Monk, K.R. (2016). Gpr126/Adgrg6 Has Schwann Cell Autonomous and Nonautonomous Functions in Peripheral Nerve Injury and Repair. J Neurosci. 36, 12351-12367.

Monk, K.R., Naylor, S.G., Glenn, T.D., Mercurio, S., Perlin, J.R., Dominguez, C., Moens, C.B., and Talbot, W.S. (2009). A G Protein-Coupled Receptor Is Essential for Schwann Cells to Initiate Myelination. Science 325, 1402-1405.

Monk, K.R., Oshima, K., Jors, S., Heller, S., and Talbot, W.S. (2011). Gpr126 is essential for peripheral nerve development and myelination in mammals. Development 138, 2673-2680.

Moriguchi, T., Haraguchi, K., Ueda, N., Okada, M., Furuya, T., and Akiyama, T. (2004). DREG, a developmentally regulated G protein-coupled receptor containing two conserved proteolytic cleavage sites. Genes to Cells 9, 549-560.

Paavola, K.J., Sidik, H., Zuchero, J.B., Eckart, M., and Talbot, W.S. (2014). Type IV collagen is an activating ligand for the adhesion G protein-coupled receptor GPR126. Sci Signal 7, ra76.

Petersen, S.C., Luo, R., Liebscher, I., Giera, S., Jeong, S.-J., Mogha, A., Ghidinelli, M., Feltri, M.L., Schöneberg, T., and Piao, X., et al. (2015). The Adhesion GPCR GPR126 Has Distinct, Domain-Dependent Functions in Schwann Cell Development Mediated by Interaction with Laminin-211. Neuron 85, 755-769.

Qin, X., Xu, L., Xia, C., Zhu, W., Sun, W., Liu, Z., Qiu, Y., and Zhu, Z. (2017). Genetic Variant of GPR126 Gene is Functionally Associated With Adolescent Idiopathic Scoliosis in Chinese Population. Spine 42, E1098-E1103.

Ravenscroft, G., Nolent, F., Rajagopalan, S., Meireles, A.M., Paavola, K.J., Gaillard, D., Alanio, E., Buckland, M., Arbuckle, S., and Krivanek, M., et al. (2015). Mutations of GPR126 are responsible for severe arthrogryposis multiplex congenita. The American Journal of Human Genetics 96, 955-961.

Schindelin, J., Arganda-Carreras, I., Frise, E., Kaynig, V., Longair, M., Pietzsch, T., Preibisch, S., Rueden, C., Saalfeld, S., and Schmid, B., et al. (2012). Fiji: an open-source platform for biological-image analysis. Nat Methods 9, 676-682.

Scholz, N., Gehring, J., Guan, C., Ljaschenko, D., Fischer, R., Lakshmanan, V., Kittel, R.J., and Langenhan, T. (2015). The adhesion GPCR latrophilin/CIRL shapes mechanosensation. Cell Rep 11, 866-874.

Scholz, N., Guan, C., Nieberler, M., Grotemeyer, A., Maiellaro, I., Gao, S., Beck, S., Pawlak, M., Sauer, M., and Asan, E., et al. (2017). Mechano-dependent signaling by Latrophilin/CIRL quenches cAMP in proprioceptive neurons. eLife 6.

Schöneberg, T., Schulz, A., Biebermann, H., Grüters, A., Grimm, T., Hübschmann, K., Filler, G., Gudermann, T., and Schultz, G. (1998). V2 vasopressin receptor dysfunction in nephrogenic diabetes insipidus caused by different molecular mechanisms. Hum. Mutat. 12, 196-205.

Slattery, A.D., Blanch, A.J., Quinton, J.S., and Gibson, C.T. (2013). Accurate measurement of Atomic Force Microscope cantilever deflection excluding tip-surface contact with application to force calibration. Ultramicroscopy 131, 46-55.

Stoveken, H.M., Hajduczok, A.G., Xu, L., and Tall, G.G. (2015). Adhesion G protein-coupled receptors are activated by exposure of a cryptic tethered agonist. Proc Natl Acad Sci U S A 112, 6194-6199.

Suchý, T., Zieschang, C., Popkova, Y., Kaczmarek, I., Weiner, J., Liebing, A.-D., Çakir, M.V., Landgraf, K., Gericke, M., and Pospisilik, J.A., et al. (2020). The repertoire of Adhesion G protein-coupled receptors in adipocytes and their functional relevance. Int J Obes (Lond).

Sun, P., He, L., Jia, K., Yue, Z., Li, S., Jin, Y., Li, Z., Siwko, S., Xue, F., and Su, J., et al. (2020). Regulation of body length and bone mass by Gpr126/Adgrg6. Science Advances 6, eaaz0368.

Takeda, K., Kou, I., Hosogane, N., Otomo, N., Yagi, M., Kaneko, S., Kono, H., Ishikawa, M., Takahashi, Y., and Ikegami, T., et al. (2019). Association of Susceptibility Genes for Adolescent Idiopathic Scoliosis and Intervertebral Disc Degeneration With Adult Spinal Deformity. Spine 44, 1623-1629.

Unal, H., Jagannathan, R., and Karnik, S.S. (2012). Mechanism of GPCR-directed autoantibodies in diseases. Advances in experimental medicine and biology 749, 187-199.

Xu, E., Lin, T., Jiang, H., Ji, Z., Shao, W., Meng, Y., Gao, R., and Zhou, X. (2019a). Asymmetric expression of GPR126 in the convex/concave side of the spine is associated with spinal skeletal malformation in adolescent idiopathic scoliosis population. European Spine Journal 28, 1977-1986.

Xu, E., Shao, W., Jiang, H., Lin, T., Gao, R., and Zhou, X. (2019b). A Genetic Variant in GPR126 Causing a Decreased Inclusion of Exon 6 Is Associated with Cartilage Development in Adolescent Idiopathic Scoliosis Population. BioMed research international 2019, 4678969.

Xu, J.-F., Yang, G.-h., Pan, X.-H., Zhang, S.-J., Zhao, C., Qiu, B.-S., Gu, H.-F., Hong, J.-F., Cao, L., and Chen, Y., et al. (2015). Association of GPR126 gene polymorphism with adolescent idiopathic scoliosis in Chinese populations. Genomics 105, 101-107.

Xu, L., Wu, Z., Xia, C., Tang, N., Cheng, J.C.Y., Qiu, Y., and Zhu, Z. (2019c). A Genetic Predictive Model Estimating the Risk of Developing Adolescent Idiopathic Scoliosis. Current Genomics 20, 246-251.

Yona, S., Lin, H.-H., Dri, P., Davies, J.Q., Hayhoe, R.P.G., Lewis, S.M., Heinsbroek, S.E.M., Brown, K.A., Perretti, M., and Hamann, J., et al. (2008). Ligation of the adhesion-GPCR EMR2 regulates human neutrophil function. FASEB J 22, 741-751.

