## Supplemental Figures for "The N terminus of adhesion G protein-coupled receptor GPR126/ADGRG6 as allosteric force integrator"

**SUPPLEMENTAL INFORMATION**

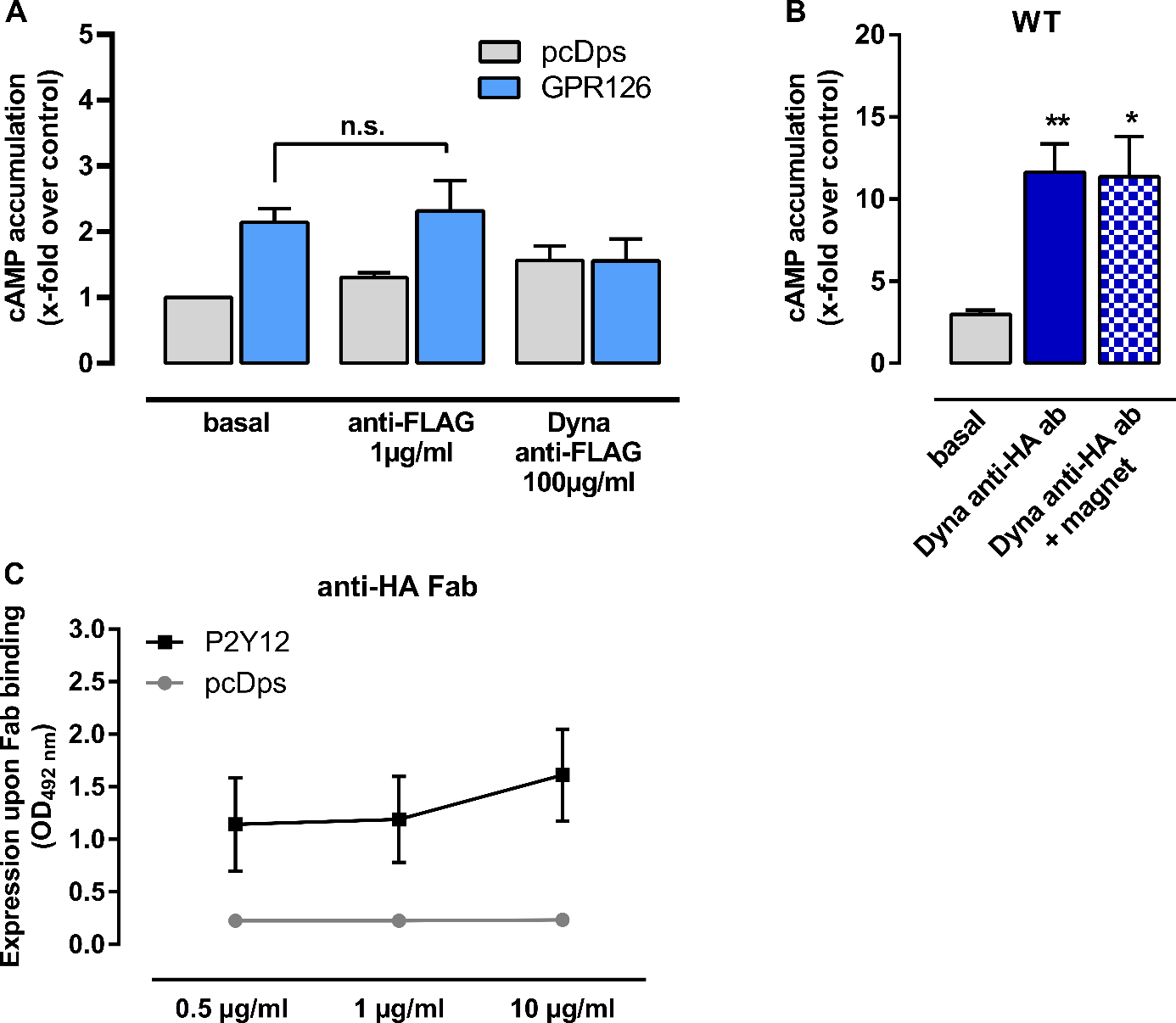

**Figure S1.** **No further activation of GPR126 can be observed using a magnet or anti FLAG antibody.** (**A**) To investigate specific activation through anti-HA antibody on GPR126, we stimulated vector control (pcDps) and GPR126-transfected COS-7 cells with anti-FLAG antibody and with anti-FLAG conjugated Dynabeads in cAMP accumulation assays. Empty vector (pcDps) served as negative control. (pcDps cAMP level: 8.96 ± 2.4 nM/well). **(B)** Cells transfected with WT GPR126 were stimulated with anti-HA ab conjugated Dynabeads® and incubated in the presence of a 700 Gs magnetic field. Compared to basal activity of the receptor a significant activation can be detected, which is, however, not different from activation induced by anti-HA ab conjugated Dynabeads® alone. Empty vector (pcDps) served as negative control (basal cAMP level in pcDps transfected cells: 7.2 ± 1.4 nM/well). (**C**) To confirm that the anti-HA Fab fragment recognizes the HA-Tag sandwich ELISA were performed. The plates were coated with the anti-HA Fab fragment. Cell lysate of cells expressing the P2Y_12_ receptor were used as sample (HA- and FLAG-tagged, positive control of all other ELISA performed) and an anti-FLAG followed by peroxidase-coupled secondary antibody was used for detection. Specific optical density (OD_492nm_) values are given. Data are given as means ± SEM of three independent experiments each performed in triplicates. Statistical analysis was performed by applying one-way ANOVA followed by Dunnett post hoc analysis; *p < 0.05; **p< 0.01.

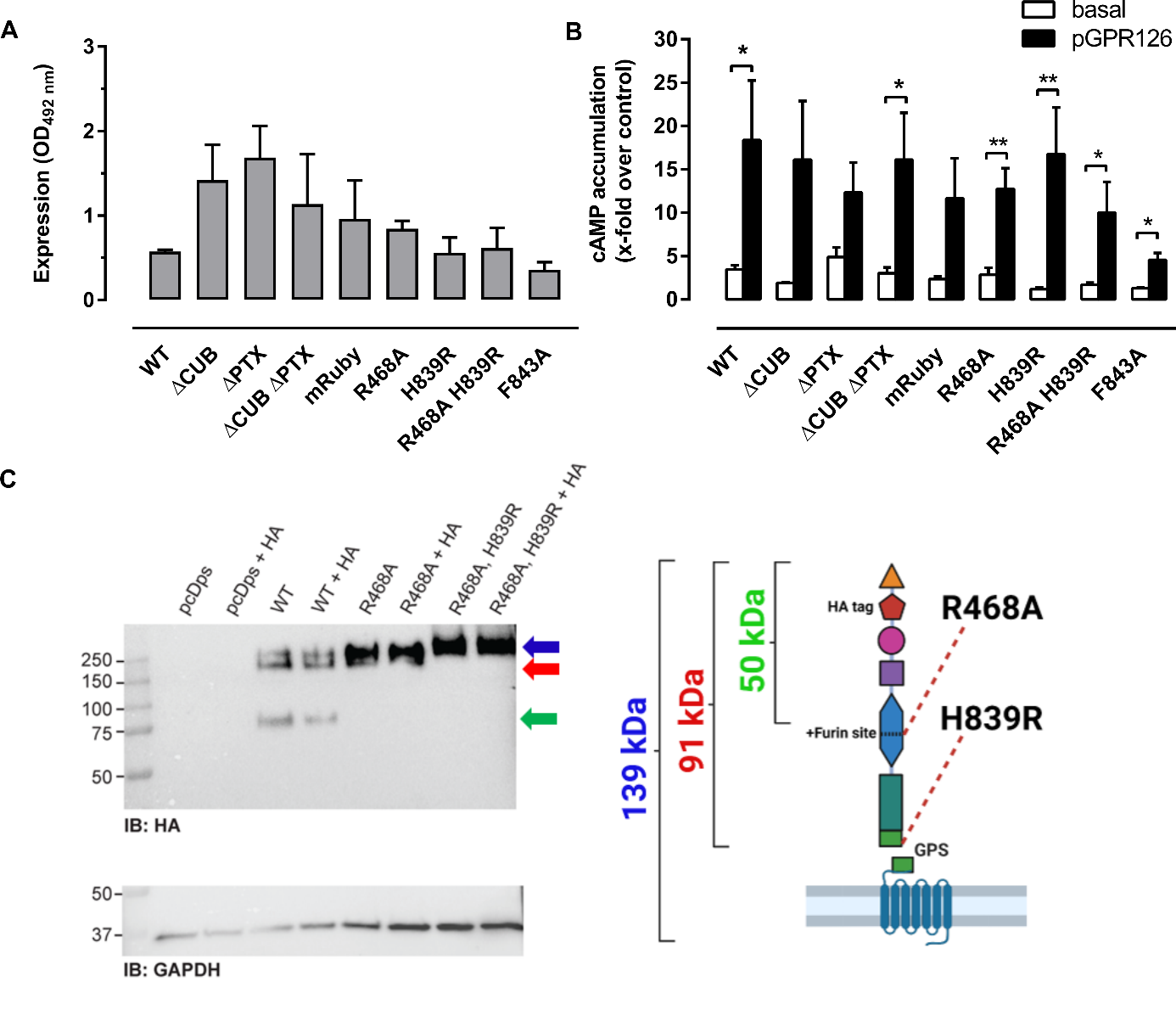

**Figure S2. Expression, activation and control of cleavage-deficiency of domain and cleavage mutants.** (**A**) For expression studies of full length and given domain mutants of GPR126 constructs in COS-7 cells, cell surface ELISA was used. Specific optical density (OD_492 nm_) readings are given. The nonspecific OD value (empty vector: 0.006 ± 0.001) was subtracted in each experiment from the specific OD value. (**B**) Wild type (WT) and given full-length GPR126 mutants were tested in cAMP accumulation assays. Receptor constructs were stimulated with 1 mM of pGPR126. Empty vector (pcDps) stimulated with 1% DMSO served as negative control (cAMP level: 4.2 ± 0.8 nM/well). Data are given as means ± SEM of at least three independent experiments each performed in triplicates. Statistical analysis was performed by unpaired t-test comparing differences between basal and peptide-stimulated activity of each mutant (*p<0.05; **p<0.01; ***p<0.001). (**C**) COS-7 were transfected with indicated constructs and 48 h after transfection cell lysates were harvested and analyzed by Western blot. Membranes were incubated with HA antibody and GAPDH antibody to detect GPR126 and the housekeeping protein, respectively. Furin cleavage is only observable in the WT construct (green arrow) and completely diminished by the R468A mutant. Molecular weight is indicated for non-glycosylated full-length receptor with 139 kDa (in blue) and a theoretical cleavage product of 50 kDa after furin-cleavage (in green) and a 91 kDa product after GPS-cleavage (in red). Corresponding bands are color coded accordingly. The unknown glycosylation pattern of the GPR126 causes a discrepancy in the Western blot bands due to additional molecular weight and changes in overall net charge.

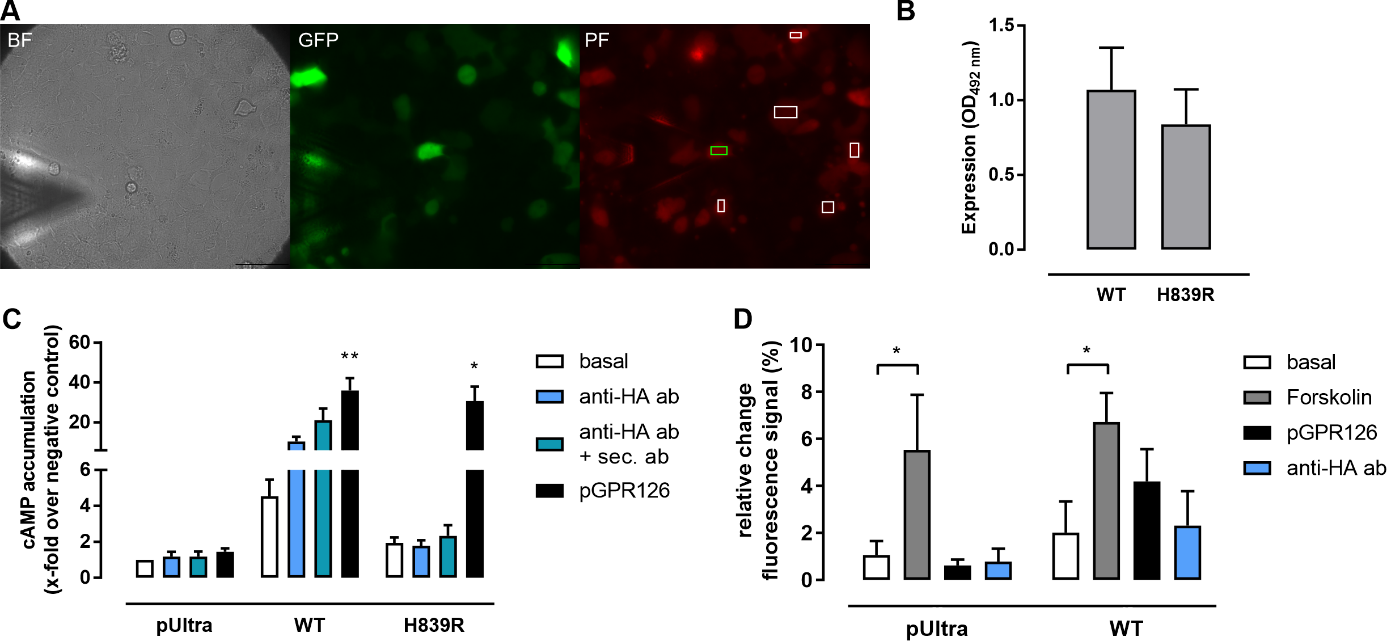

**Figure S3. Evaluation of AFM images, expression of WT and cleavage mutant H839R and their activation in HEK-GT cells and mechano-independent changes in the Pink Flamindo fluorescence signal. (A)** Imaging of the successfully co-transfected cell with the AFM cantilever positioned in the bottom left corner: Brightfield (BF) and Green Fluorescent Protein (GFP). Pink Flamindo (PF) images were acquired after the AFM tip was moved onto the stimulated cell right before and right after pressure or pulling forces were applied. Fluorescence intensity changes were measured between these two images. The green rectangle marks the area in which the brightness of the stimulated cell was measured. The white rectangles mark the areas used as reference to account for changes in fluorescence intensity happening independently from the AFM cantilever stimulation. (**B**) For expression studies of the full length and the cleavage-deficient GPR126 mutant in the pUltra vector in HEK-GT cells, cell surface ELISA was used. Specific optical density (OD_492 nm_) readings are given. The nonspecific OD value for empty vector was 0.008 ± 0.001. (**C**) The WT and the cleavage-deficient full-length GPR126 constructs were tested in cAMP accumulation assays. Receptor constructs were stimulated with 1 mM of pGPR126, anti-HA ab (1 µg/ml) and anti-HA ab (1 µg/ml) + secondary ab (22 µg/ml). Empty vector (pUltra) served as negative control (stimulated with 1% DMSO as a negative control for pGPR126 stimulation) (basal: cAMP level: 2.54 ± 0.45 nM/well, 1% DMSO cAMP level: 2.48 ± 0.51 nM/well). Data are given as means ± SEM of three (ELISA) or six (cAMP accumulation assay) independent experiments each performed in triplicates. Statistics were performed by applying one-way ANOVA followed by Dunnett post hoc analysis; *p < 0.05; **p < 0.01. All significances given in the graph show the result of the post hoc analysis and compare the condition to the basal cAMP accumulation of the same construct. **(D)** Cells co-transfected with Pink Flamindo and either pUltra or GPR126 pUltra DNA were stimulated with forskolin (10 µM), pGPR126 (1 mM) or anti-HA ab (1 µg/ml) and the changes in Pink Flamindo fluorescence intensity occurring after 60s were measured. Statistics were performed by applying one-way ANOVA followed by Dunnett post hoc analysis; *p < 0.05. Significances given in the graph show the results of the post hoc analysis.
